# Can offsetting the energetic cost of hibernation restore an active season phenotype in grizzly bears (*Ursus arctos horribilis*)?

**DOI:** 10.1101/2021.03.12.435162

**Authors:** Heiko T. Jansen, Brandon Evans Hutzenbiler, Hannah R. Hapner, Madeline L. McPhee, Anthony M. Carnahan, Joanna L. Kelley, Michael W. Saxton, Charles T. Robbins

**Author notes:** **Correspondence:** Heiko T. Jansen, PhD. **Conflict of Interest:** The authors declare that the research was conducted in the absence of any commercial or financial relationships that could be construed as a potential conflict of interest. **Author Contributions:** HTJ, CTR and JLK obtained funding; HTJ, CTR and BEH designed the study; HTJ, CTR, BEH, HRH, AMC, JLK and MWS performed the oral glucose tolerance tests, blood and tissue sampling; BEH, MLM and HRH performed the metabolic flux analyses and serum assays; HTJ, BEH and HRH analyzed the data; all authors contributed to the writing and editing of the manuscript. HTJ approved the final manuscript version for submission. **Funding:** Funding was obtained from the International Association for Bear Research and Management and the following Washington State University endowments: Raili Korkka Brown Bear Fund, Bear Research and Conservation Fund and Nutritional Ecology Fund.

## Abstract

Hibernation is characterized by suppression of many physiological processes. To determine if this state is reversible in a non-food caching species, we fed hibernating grizzly bears (*Ursus arctos horribilis*) glucose for 10 days to replace 53% or 100% of the estimated minimum daily energetic cost of hibernation. Feeding caused serum concentrations of glycerol and ketones (ß-hydroxybutyrate) to return to active season levels irrespective of the amount of glucose fed. By contrast, free-fatty acids and indices of metabolic rate, such as general activity, heart rate, and strength of the daily heart rate rhythm and insulin sensitivity were restored to roughly 50% of active season levels. Body temperature was unaffected by feeding. To determine the contribution of adipose to these metabolic effects of glucose feeding we cultured bear adipocytes collected at the beginning and end of the feeding and performed metabolic flux analysis. We found a roughly 33% increase in energy metabolism after feeding. Moreover, basal metabolism before feeding was 40% lower in hibernation cells compared to fed cells or active cells cultured at 37°C, thereby confirming the temperature independence of metabolic rate. The partial suppression of circulating FFA with feeding likely explains the incomplete restoration of insulin sensitivity and other metabolic parameters in hibernating bears. Further suppression of metabolic function is likely an active process. Together, the results provide a highly controlled model to examine the relationship between nutrient availability and metabolism on the hibernation phenotype in bears.

## INTRODUCTION

The ability to hibernate or express torpor (a shorter period of metabolic suppression and body temperature reduction) may have evolved multiple times and suggests that different cellular mechanisms can produce this phenotype (Geiser, 1998; Geiser, 2004). Hibernation in rodents has long captured the attention of physiologists due to the extreme changes that occur seasonally in metabolism, body temperature, and body mass (Carey et al., 2003; Mohr et al., 2020; Nelson et al., 1983; Storey and Storey, 1990). However, hibernating brown bears (*Ursus arctos*) exhibit a very different phenotype from what occurs during rodent hibernation (Harlow et al., 2002; Hellgren, 1998; Jansen et al., 2016; Lane et al., 2012; Lin et al., 2012; Lohuis et al., 2005; Nelson et al., 1983; Robbins et al., 2012; Toien et al., 2011; Ware et al., 2012). Thus, comparative studies provide an ideal opportunity to explore the different genetic, physiological, biochemical and environmental underpinnings of hibernation.

For virtually all hibernators climate change and other anthropogenic factors present new challenges, and it is important to determine if these factors could alter, or even eliminate, the expression of this ancestral phenotype (Geiser, 1998; Geiser, 2013; Lane et al., 2012). One extensively studied factor promoting hibernation is the amount and quality of food (Florant and Healy, 2012; Frank et al., 2008; Harlow and Frank, 2001; Siutz et al., 2017; Vuarin and Henry, 2014). Bears and many other seasonal hibernators undergo extreme annual mass gains due to increasing adiposity prior to hibernation (Carey et al., 2003; Dark, 2005). Along with this high level of adiposity preceding the onset of hibernation, hibernating bears reduce their activity levels by more than 90% (Robbins et al., 2012; Ware et al., 2012), develop insulin resistance (Kamine et al., 2012a; McCain et al., 2013; Palumbo et al., 1983; Rigano et al., 2017), and reduce their heart rate by as much as 90% relative to the active state while maintaining a relative high body temperature in comparison to rodents (Laske et al., 2017; Nelson et al., 2010; Toien et al., 2011).

Although the impact of food availability on torpor in heterothermic endotherms has been extensively studied (see Review by Vuarin and Henry, 2014), several aspects such as the lack of discrete torpor-arousal bouts, large body mass, and birth of young in the den, are unique to bears. Thus, to further explore the genetic and physiological controls of bear hibernation, we sought to develop an experimental paradigm whereby some or all aspects of hibernation could be reversed. To this end we reasoned that by feeding a single nutrient, glucose, rather than a complex diet, we could relate energy supply to energy demand via alterations in metabolic profiles and physiology during hibernation. Similarly, if the cellular changes that are triggered by consuming a single nutrient were preserved in cells from a critical tissue such as adipose, we then would have a robust system of three metabolic states (e.g., hibernation, fed-hibernation, and fed-active) to explore many basic aspects of hibernation both in vivo and in vitro. The present study was therefore performed to test the hypothesis that glucose feeding is capable of reversing the hibernation phenotype in bears.

## METHODS

### Animals

Grizzly bears (*Ursus arctos horribilis*, Linnaeus 1758, n=11) were housed at the Washington State University Bear Research, Education and Conservation Center (WSU Bear Center, Pullman, Washington, USA) in accordance with the Bear Care and Colony Health Standard Operating Procedures approved by the Washington State Institutional Animal Care and Use Committee based on U.S. Department of Agriculture guidelines and in accordance with current animal care and use guidelines approved by the American Society of Mammalogists (Sikes et al., 2011), protocols #04952, #06546 and #06468. The bears in the facility hibernate from November to mid-to-late March. Bears of both sexes were used with ages ranging from 3-15 years when the study began. Diet and feeding schedules during April-October (active season) were similar to those described previously (Rigano et al., 2017). Bears were monitored continuously using video cameras mounted in each den, outdoor run and outdoor exercise yard.

### Feeding during hibernation

Bears at the WSU Bear Center are trained for blood sampling year-round using honey diluted 12.5-25% with water (v:v) (Joyce-Zuniga et al., 2016; Ware et al., 2013). Bears are highly motivated to receive honey during blood sampling based on positive-reinforcement training and can be sampled without the use of sedatives or other drugs even during hibernation (Joyce-Zuniga et al., 2016). Thus, feeding glucose (dextrose, Sigma-Aldrich, St. Louis, MO) during hibernation is a simple extension of this approach. The amount of dextrose fed was calculated to replace the estimated daily energy lost based on the least observed metabolic rate (LOMR) of hibernating grizzly bears using equation 1.

*1) Y=4*.*8X*^*1*.*09*^ (Robbins et al., 2012)

*where Y is least observed metabolic rate (kcal/day) and X is body weight (kg*)

We chose two levels of energy replacement based on the LOMR: 53% (n=7) and 100% (n=6). The 53% level was fed in January (mid-hibernation) 2017-2018 and corresponds to a 1g/kg^-1^day^-1^ glucose which is the standard dose used for our oral glucose tolerance tests oGTT (see below) and used to confirm the insulin resistance of hibernating bears (Rigano et al., 2017). The 100% level (range 1.82-1.93 gkg^-1^ depending on bear weight) was fed in January 2018-2019. Four bears served as unfed controls. For both feeding trials, glucose was diluted in water to the same concentration, warmed to approximately 22°C and fed once daily at 9AM for 10 consecutive days. The volume of dextrose fed daily was based on the mass measured at the start of hibernation minus the predicted mass loss that occurred since the beginning of hibernation (Robbins et al., 2012). The 10-day feeding duration was chosen to allow for comparisons of glucose disposal (see below) and for sufficient recovery of energetic and metabolic parameters without obscuring hibernation altogether. The timeline for feeding and related procedures is shown in Fig. S1.

### Tissue and blood sampling

Fat biopsies were obtained using a 6mm biopsy punch after bears were anesthetized with a combination of tiletamine HCl/zolazepam HCl (Telazol®; Zoetis, Florham Park, NJ) and dexmedetomidine (Zoetis) as described previously (Rigano et al., 2017). At this time larger volumes of blood samples were also collected (∼150ml) from the jugular vein into 10ml Tiger-Top tubes (BD Vacutainer SST tubes) for use in cell culture experiments. Blood was allowed to clot, centrifuged within 3-4h, aliquoted and the serum was frozen at −80°C until assayed or used for cell culture (see below). Serum included: hibernation (pre-feeding; HIBS), serum from fed bears (DEXS), serum from active season bears (ACTS). Glucose determinations were made in the trained unanesthetized bears (fed group only) in 1 cc syringes while individual bears were briefly housed in a movable crate described previously (Joyce-Zuniga et al., 2016; Rigano et al., 2017).

### Oral glucose tolerance tests (oGTT)

Bears in the fed group were administered an oGTT using a standard 1gkg-1 dose of glucose to infer the relative state of insulin sensitivity as previously described (Rigano et al., 2017). All oGTT were performed at the end of the 10-day feeding trial only in unanesthetized (fed) bears to avoid excessive glucose exposure and the confounding influence of anesthetics on glucose disposal and other physiological parameters (Kamine et al., 2012b; Nelson and Robbins, 2010). To track the ongoing changes in insulin sensitivity during feeding periods we collected blood samples from a peripheral metatarsal vein before (0min) and 120min after glucose feeding from day 0 (pre-feeding, i.e., hibernation) until day nine (Fig. S1). Since our bears are trained with honey for blood sampling without the use of anesthesia and honey feeding would confound the blood glucose determinations following glucose administration, we used a non-caloric substitute (Stevia extract, 8ml diluted in 500ml water) to collect the second (i.e., 120min) blood sample during the 10-day feeding procedure and for the second and subsequent (15, 30, 60, 120 min) blood samples collected during the oGTT. The amount of Stevia extract was kept to a minimum in an effort to reduce volume effects. All blood glucose determinations were made in duplicate immediately after collection using a calibrated Accu-Chek Aviva glucometer (Roche, Basel, Switzerland).

### Serum metabolites, insulin and glucagon

Serum collected at the time of biopsy before feeding began (i.e., mid hibernation) and at the time of the second biopsy was assayed for glycerol, free fatty acids (FFA) and ketones (ß-hydroxybutyrate) using commercial kits (Cayman Chemical, Ann Arbor, MI) according to the manufacturer’s instructions. All samples were run in duplicate and corrected for assay blanks where appropriate. Serum insulin and glucagon were determined using a commercial porcine/canine ELISA (Alpco, Salem, NH) and multi-species EIA (Phoenix, Burlingame, CA), respectively, as described previously (Rigano et al., 2017).

### Heart rate and core body temperature

As seasonal changes in heart rate closely follow those of metabolic rate in bears, heart rate can serve as a useful proxy (Nelson and Robbins, 2010; Toien et al., 2011). We used five bears (4 fed and 1 unfed) and implanted them with small cardiac monitors developed for human heart patients (Reveal LINQ, Medtronic, Minneapolis, MN; 4.0 mm X 7.2 mm X 44.8 mm; 2.4 grams) which were capable of recording heart rate at 2 min intervals and body temperature at 4h intervals continuously for up to two years. For implantation bears were anesthetized as described above with a combination of Telazol and dexmedetomidine and surgically prepared using standard procedures. Devices were implanted subcutaneously in left peristernal locations with surgical sutures used to close the puncture sites. All bears were monitored closely for signs of irritation and/or rejection. Data from Dec 2017 to late Feb 2018 (53% feeding) and from Nov 2018 to Mar 2019 (100% feeding) were analyzed for this study as described below.

### Heart rate and body temperature analysis

Heart rate data were analyzed to determine the effect of feeding on overall metabolic status by comparing mean values for the 10 days prior to feeding (and before biopsy), the 10 days of feeding and for the 10 days after feeding ended (also after biopsy and 5-day recovery).

Additionally, to examine the impact of feeding on daily (circadian) heart rate rhythms, we quantified the strength of the daily heart rate rhythm (%) and its period (hr) using custom Matlab scripts described previously (Jansen et al., 2016). Circadian rhythm strength is defined as the proportion of variance (range 0-100) in the 12-64 h period frequency band and was determined from the discrete wavelet transforms (Jansen et al., 2016). Body temperature data were not analyzed for rhythm strength or period given the long (4h) sampling interval. Only 10-day mean body temperature data were analyzed for feeding effects.

### Activity determinations

General movements (hereafter referred to as ‘activity’) were scored manually daily during the 10 days prior to feeding, during the 10-day feeding period and then for 50 days after glucose feeding using video recordings of each den. Four, 1h blocks (between 0700-0800h, 1100-1200h, 1500-1600h and 2300-2400/0000h) were analyzed each day. Each hourly block was then divided into 6, 10min epochs. For each epoch an observer recorded an ‘activity’ bout if a bear stood on all four legs, walked, sat up or reared on its hind legs. Then, the proportion of the 1h block each bear spent active was calculated. The average percent time spent standing for each hourly epoch for all fed and unfed bears was then calculated and analyzed over course of the study to allow for statistical analysis of daily and long-term trends.

### Adipocyte cell culture

Mesenchymal stem cells from grizzly bears were obtained from subcutaneous gluteal fat using 6mm punch biopsies during late May (active season; ACT), early January (pre fed; HIB) and late January (post fed, dextrose; DEX). Samples were enzymatically dissociated using Liberase-TM (Sigma-Aldrich) to obtain the stromal vascular fraction (SVF), plated in 12-well culture plates, expanded and cryopreserved as described by Gehring (Gehring et al., 2016). For oxygen consumption and glucose uptake experiments, cryopreserved cells were thawed, seeded for culture (2500-7000 cells per well) in Seahorse XFp Miniplates (Agilent, Santa Clara, CA, see below) or 96-well culture plates and then differentiated into mature adipocytes according to minor modifications of our previously published protocols (Gehring et al., 2016; Rigano et al., 2017). Briefly, SVF cells were grown in 89% low glucose (5.55mM) DMEM (ThermoFisher, Waltham, MA) containing glutaMAX, 10% fetal bovine serum (FBS; Atlanta Biological now Bio-Techne, Minneapolis, MN) and 1% PSA antibiotic/antimycotic (100 units ml^-1^ penicillin, 100µg ml^-1^ streptomycin and 0.25 µg ml^-1^ amphotericin B) until approximately 80-90% confluent - usually 2-3 days. Differentiation into mature adipocytes was induced with medium containing DMEM, 1% PSA, 10% serum (either bear active (ACTS), hibernation (HIBS), or post fed hibernation (DEXS) serum pooled from individual bears 1:1) or FBS, 861nM insulin, 1nM triiodothyronine (T3), 0.5mM IBMX, 1.1µM dexamethasone, 0.5 µM rosiglitazone and 125 µM indomethacin for two days, producing changes associated with adipogenesis (Gehring et al., 2016). Differentiation medium was then removed and replaced with a maintenance medium containing low glucose (5.5mM) DMEM, 1% PSA, 10% bear serum or FBS, 861nM insulin, 1nM T3 and 0.5 µM rosiglitazone for two days, followed by the same medium composition with 1.0 µM rosiglitazone for an additional four days. Cells were assayed eight days post induction of differentiation.

### Cellular Glucose Uptake

We quantified glucose uptake by measuring the difference in medium glucose concentrations before and after insulin stimulation using a standard glucometer. The same protocol was followed to grow cells in 96-well culture plates, as described above. In these experiments, low glucose DMEM containing 1% PSA and +/- insulin (1000nM) was applied for 12h to cells grown for 8 days post induction of differentiation in different serum conditions described above Prior to the insulin application, cells were washed 1x with PBS and cultured in low glucose DMEM w/ 1% PSA without serum overnight.

### Oxygen consumption and glycolysis determinations

Cryopreserved cells from hibernating, fed and active season bears were thawed and plated in 8-well Seahorse XFp Miniplates (Agilent) and processed as described above. Phenotype tests were carried out as described by the manufacturer’s instructions, with minor modifications to optimize our protocol. On the day of metabolic measurements, the cell culture medium was removed, the cells were washed twice and replaced with assay medium (Seahorse Base Media (103193-100) containing 5.5mM glucose, 4mM glutamine and 2mM pyruvate, pH 7.4). Plates and assay medium were then placed into a 37C incubator without CO_2_ for 60 min to allow for pH stabilization and outgassing (Pike Winer and Wu, 2014). During this time, the sensor cartridges were loaded with a combination of 2uM oligomycin and 1uM FCCP to assess the cellular phenotype and placed in Seahorse XFp analyzer (Agilent) to be equilibrated/calibrated prior to the assay run. Then, just before loading the miniplates, one final medium change was performed using outgassed medium. At the completion of the preparatory steps, the XFp cell Miniplate was loaded into the XFp analyzer for the determination of mitochondrial respiration based on oxygen consumption rates (OCR) and glycolytic flux based on extracellular acidification rates (ECAR) (Pike Winer and Wu, 2014). Basal ECAR (mpH/min) and OCR (pmol/min) were determined in six cycles defined by a two-minute mixing, zero-minute wait and a two-minute measure for a total duration of 24 minutes. After the sixth basal read, a stress mixture containing 2uM oligomycin and 1uM FCCP was injected into each well. Stressed ECAR and OCR were then determined using the same cycle parameters as basal conditions (total assay duration 48 minutes). All experiments were performed at 37°C as the operating temperature of our XFp analyzer was not adjustable. Total protein was determined for each well at the completion of each experiment using a Pierce BCA assay; all rates are reported per min/per µg protein. Values represent the average across seven individual bears, with 2-4 technical replicates (n) per bear per treatment (HIB FBS (3), HIB ACTS (2), HIB DEXS (3), HIB HIBS (4), ACT FBS (2), ACT ACTS (4)).

### Statistical analysis

All data were analyzed for treatment and time course effects using Prism v.8.0 (Graphpad Software, San Diego, CA). One- or two-way ANOVA or mixed effects models were used to compare pre-feeding, feeding and post-feeding data. OCAR and ECAR results were compared using one-way ANOVA of normalized data. Metabolite data were analyzed using paired t-test. Post-hoc analyses were performed (where appropriate) using Holm-Šidák correction for multiple comparisons. Bear, Bear x Day (or time) and Bear x Feeding level were used a random effect and Geisser-Greenhouse correction was applied in all mixed effects models.

## RESULTS

### 1. Glucose utilization

#### a. oGTT

The characteristic elevation of blood glucose at 120 min following an oral glucose challenge and indicative of insulin resistance was partially reversed by feeding glucose during hibernation, F(1,6)=121.8, P<0.0001. Once feeding began, the blood glucose at 120 min began a progressive decline over the ten days (Fig. 1). The decline was less pronounced in the 100% feeding group (main effect of feeding level – F(1,6)=13.35, P=0.0107). However, by day nine the 120 min blood glucose concentrations were similar between 53% and 100% fed groups and reached an intermediate concentration relative to the pre-feeding 0- and 120-min values (Fig. 1). oGTT performed at the conclusion of the ten-day feeding period confirmed the increase in glucose disposal. However, there were no significant differences between feeding levels (Fig. 2A; main effect of feeding level – F(1,39)=0.102, P=0.751). When compared to active season glucose profiles from June 2019, the blood concentrations at 120 min from dextrose-fed bears were not different from active season values but were significantly lower than pre-feeding hibernation levels in 2019 (t(18)=3.385, P=0.0196; Blue line in Fig. 2A).

**FIGURE 1.**
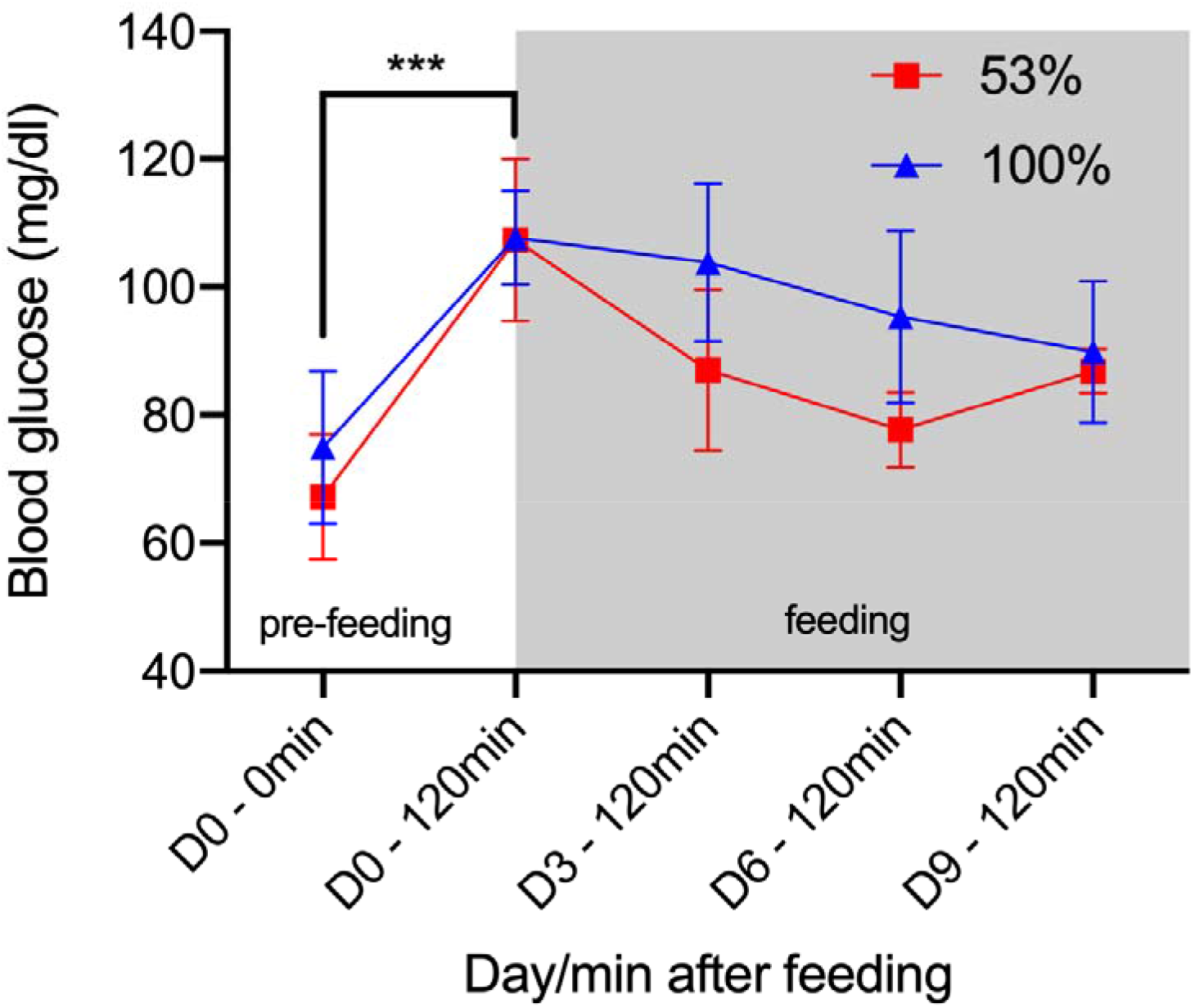
Blood glucose concentrations (mean ±SD) in trained (unanesthetized) hibernating bears before feeding glucose (day 0) and during feeding on days 3, 6, and 9. Only 120 min samples were collected from trained bears during feeding to minimize any effect of disturbance.

**FIGURE 2.**
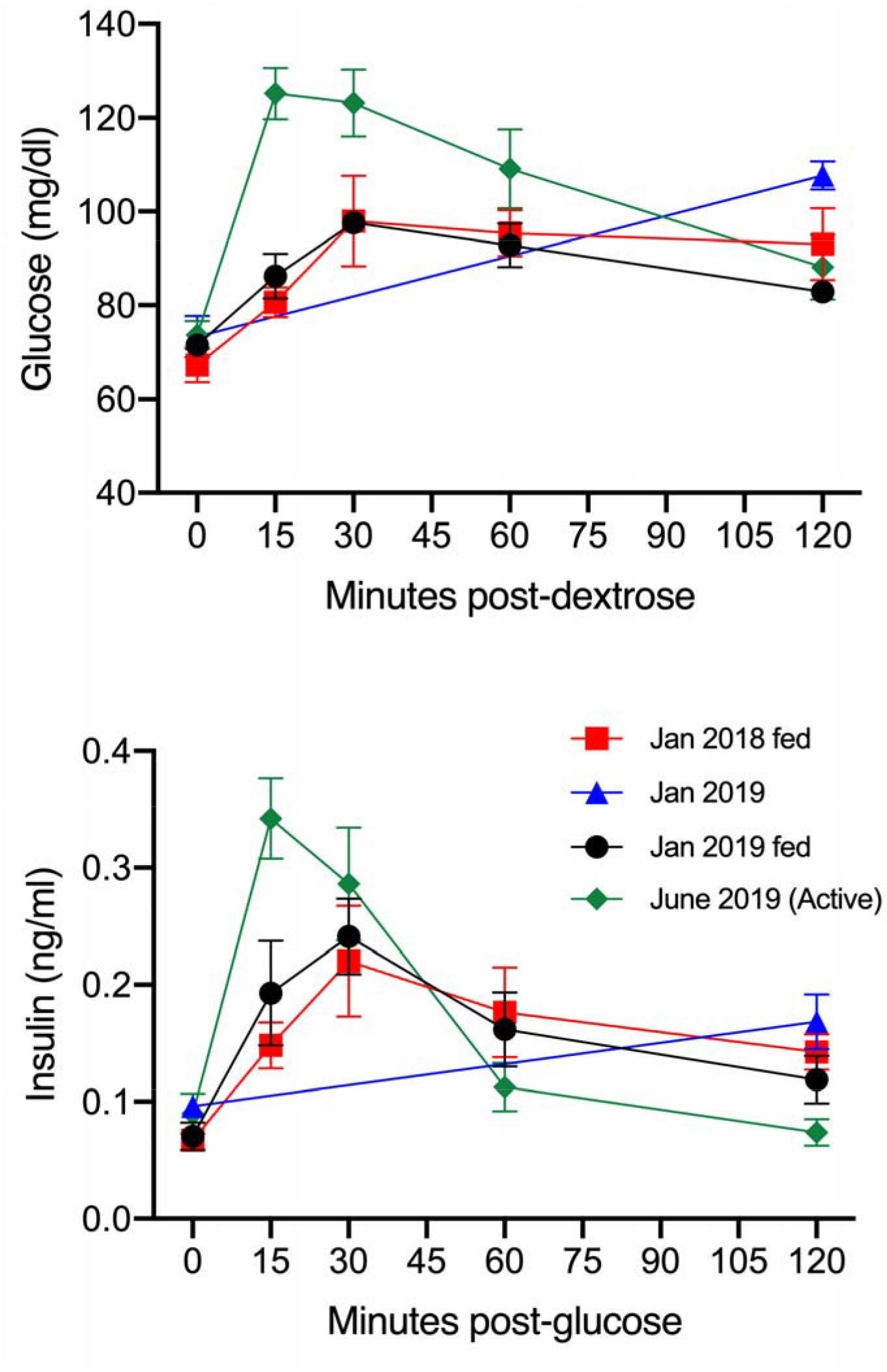
Mean±SEM blood glucose and insulin concentrations prior to (0 min) and during an oral glucose tolerance test (1g/kg glucose). Active and hibernation seasons are shown for comparison. The pre-feeding hibernation glucose data shown in blue are the same as shown in Fig. 1.

Serum insulin exhibited similar trends to blood glucose prior to feeding (Fig. 2B). We found no significant differences in insulin profiles between 53% and 100% feeding levels (F(1,31)=0.096, P=0.793). However, these profiles differed significantly from those in June 2019 (active season; time x season interaction – F(8,62)=3.056, P=0.0058).

#### b. Cellular glucose uptake

An overall effect of insulin (F(1,48)=22.39, P<0.0001), serum (F(3,48)=11.09, P<0.0001 and interaction was revealed (F(3,48)=5.019, P=0.0042) in hibernation adipocytes (Fig.3 HIB CELLS). However, post-hoc analysis revealed that the enhanced insulin response was due solely to cells cultured in active season serum (ACTS; t(48)=5.658, P<0.0001). Cells from fed bears (Fig. 3 DEX CELLS) also exhibited an overall effect of insulin-stimulated glucose uptake (F(1,48)=72.27, P<0.0001), serum (F(3,48)=6.115, P=0.0013 and interaction (F(3,48)=3.077,

**FIGURE 3.**
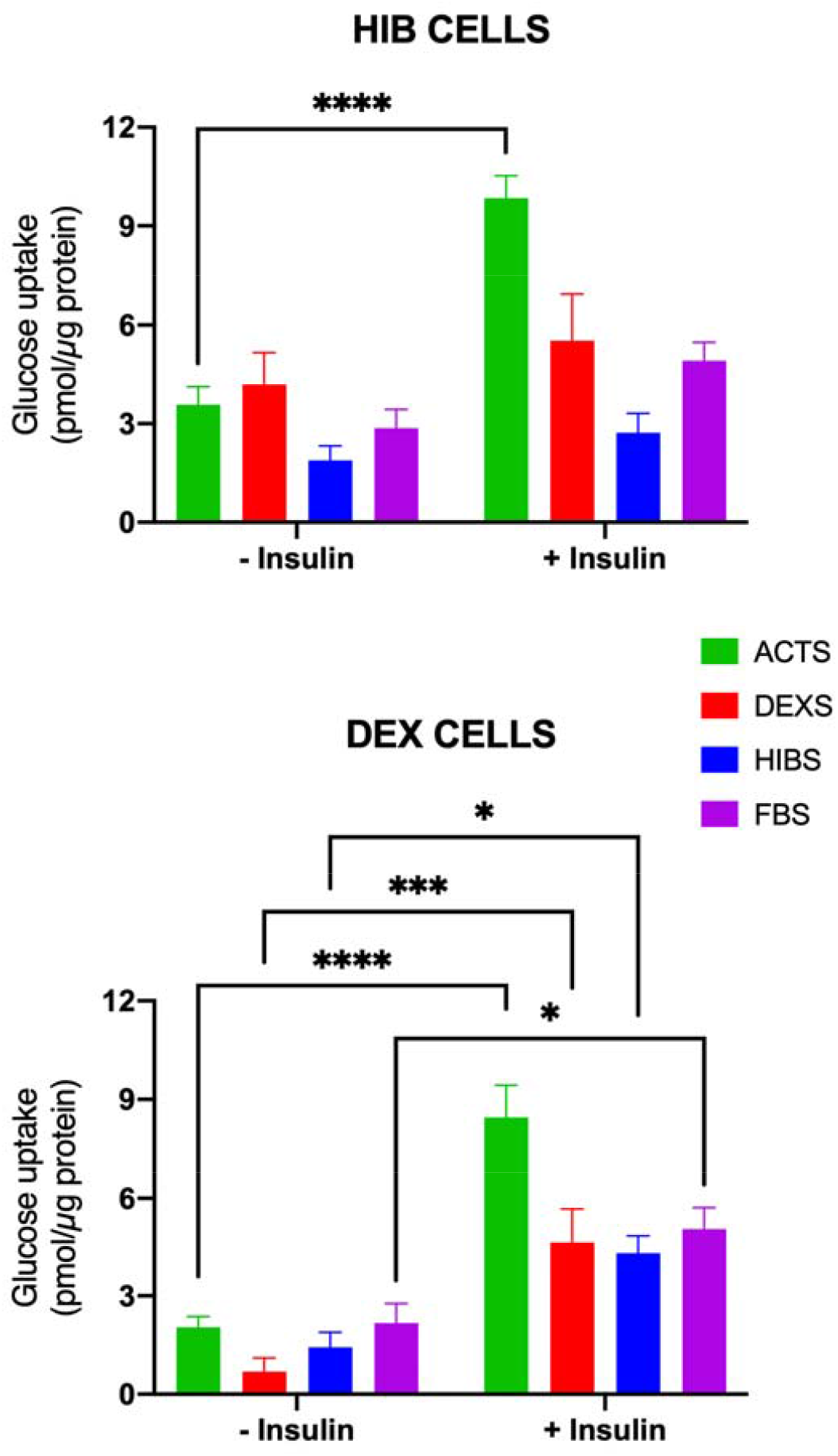
Twelve-hour glucose uptake (mean ± SEM) by bear adipocytes in hibernation (pre-feeding, HIB CELLS) and cells from fed bears (DEX CELLS) without or with insulin (1000nM) and under different serum conditions (ACTS-active season serum; DEXS-fed serum (53% fed); HIBS-hibernation serum or FBS). *P<0.05, ***P<0.001, ****P<0.0001.

P=0.0362). In contrast to hibernation cells, cells from fed bears responded to insulin under all serum conditions. The effect (fold-change from baseline) was greatest in cells incubated in serum from fed bears (DEXS; approximately 6.5-fold); this response was greater than in fed cells cultured in ACTS (approximately 4-fold) or hibernation cells cultured in ACTS (approximately 3-fold).

### 2. Serum indices of metabolic status

Daily glucose feeding for 10 days resulted in significant decreases in serum glycerol, FFA and ß-hydroxybutyrate concentrations compared to pre-feeding levels (Table 1). However, these reductions did not differ with level of glucose feeding (glycerol – F(1,10)=0.5605, P=0.2400; FFA – F(1,10)=0.9270, P=0.3583; ß-hydroxybutyrate – F(1,10)=0.3786, P=0.5521). Pre-feeding levels were similar to those in unfed bears while post-feeding levels were similar to those in the active season (Table 1). Glucagon concentrations were unaffected by feeding (not shown).

**Table 1.** Impact of glucose feeding on mean (±SEM) serum concentrations of lipolysis products and ketones in hibernating bears (n=6)^†^. Bears were fed glucose to replace 53% or 100% of the predicted cost of hibernation (see Methods for details). Data from unfed hibernating bears (n=4; 2019) and fed active season bears (n=11) are shown for comparison.

| | Glycerol (μmol/L) | | FFA (μmol/L) | | β-hydroxybutyrate (μmol/L) | |
| --- | --- | --- | --- | --- | --- | --- |
|  | 53% | 100% | 53% | 100% | 53% | 100% |
| Pre-feeding | 122.38<br>(15.81) | 155.49<br>(20.31) | 376.47<br>(22.92) | 446.93<br>(43.87) | 507.83<br>(10.82) | 510.67<br>(19.07) |
| Post-feeding | 65.69<sup>b</sup><br>(10.21) | 57.65<sup>b</sup><br>(13.25) | 167.2<sup>a</sup><br>(5.48) | 178.86<sup>a</sup><br>(25.93) | 183.67<sup>b</sup><br>(15.96) | 162.71<sup>b</sup><br>(28.71) |
| Unfed | 109.24 (19.65) |  | 346.81 (80.26) |  | 437.13 (51.4) |  |
| Active | 48.86 (15.09) |  | 82.02 (24.01)* |  | 131 (10.6) |  |

a - P<0001 vs. Pre-feeding
b - P<0.05 vs Pre-feeding
\* - different from Post-feeding, P<0.01
<sup>†</sup> - One bear was removed from the study in 2019 (100% group).

### 3. General activity

Very low levels of activity were observed in all hibernating bears prior to the beginning of feeding (Fig. 4). A blunted, yet clearly evident daily rhythm of activity was present in fed and unfed bears (Fig. 4C,D). Since hourly pre-feeding data were only available for 1500h in the 53% group (2017-2018), direct comparisons between glucose groups were not possible. Nevertheless, activity levels were at their lowest in both groups of bears prior to the beginning of feeding. Daily glucose feeding resulted in significant increases in activity at 0700, 1200 and 1500h (Fig. 4). The effect of feeding on activity was evident for up to 50 days post feeding in both 53% and 100% groups. However, at 50 days post-feeding, the increase in activity coincided with the natural increase in activity as seen in the unfed bears prior to the end of hibernation. The increase was still nearly twice that of the unfed bears.

**FIGURE 4.**
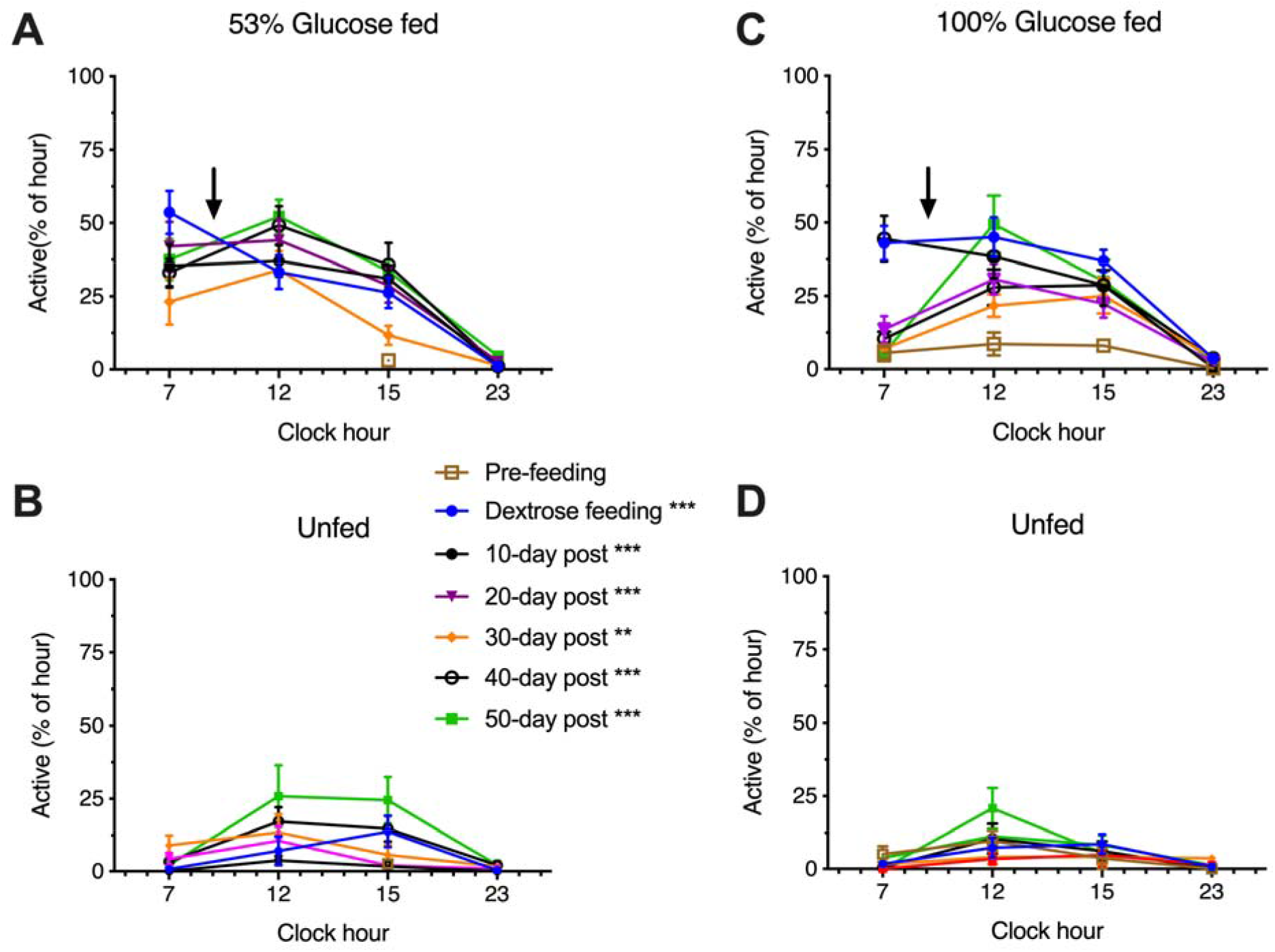
Daily mean (±SEM) activity (% of each hour standing) in 10-day blocks for fed (A,C) and unfed bears (B,D). Arrows indicate time of feeding (9AM) during the 10-day dextrose feeding phase. ** P<0.01 vs. unfed; *** P<0.001 vs. unfed. n=7 (53%), n=6 (100%). P-values are the same for 53% and 100% feeding groups.

### 4. Heart rate and body temperature

The heart rate of hibernating bears prior to any manipulations ranged from 10 to 13 beats per min (bpm) in fed bears (Fig. 5). Upon feeding, heart rate increased significantly (F(1,12)=5.101; P=0.043); however, no significant difference between feeding levels was found (F(1,12)=0.002; P=0.962). Heart rate was elevated in both 53% and 100% groups for the duration of monitoring or until hibernation ended (Fig. 6). By contrast, heart rate of the unfed bear remained low until March when it began a progressive increase (Fig. 6, 100%). A similar increasing trend was observed for all of the fed bears. All bears exhibited an increase in heart rate at the time of biopsy (arrows, Fig. 6), but this returned to roughly pre-biopsy levels within about five days (i.e. during the recovery period). We observed several transient heart rate excursions in the unfed bear (e.g., on 1/11/2018, Fig. 6, 53%) when other bears were being fed. These transients were likely due to brief disturbance as all bears were housed in the same facility, but in different pens. Heart rate returned to low, hibernation levels in the unfed bear once feeding of the other bears ended (Fig. 6A).

**FIGURE 5.**
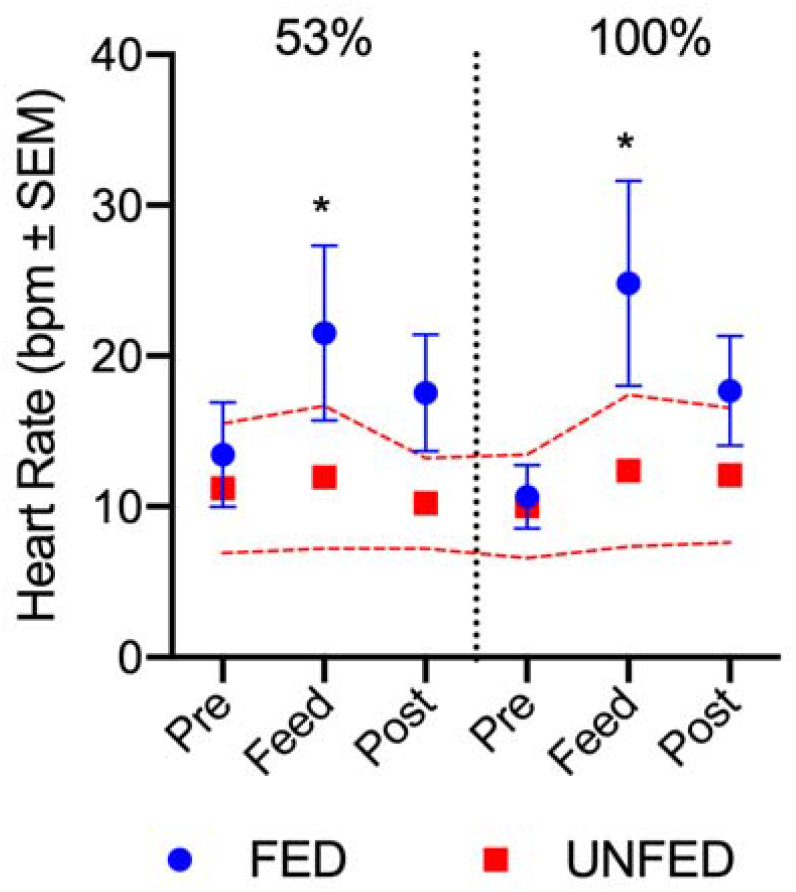
Mean heart rates in bears (N=4) fed two different levels of energy replacement. The red dashed line represents the 95% confidence interval for a single unfed bear. Each point represents a 10-day average collected prior to feeding (Pre), during feeding (Feed), and after feeding stopped (Post). * P<0.05 vs. Pre.

**FIGURE 6.**
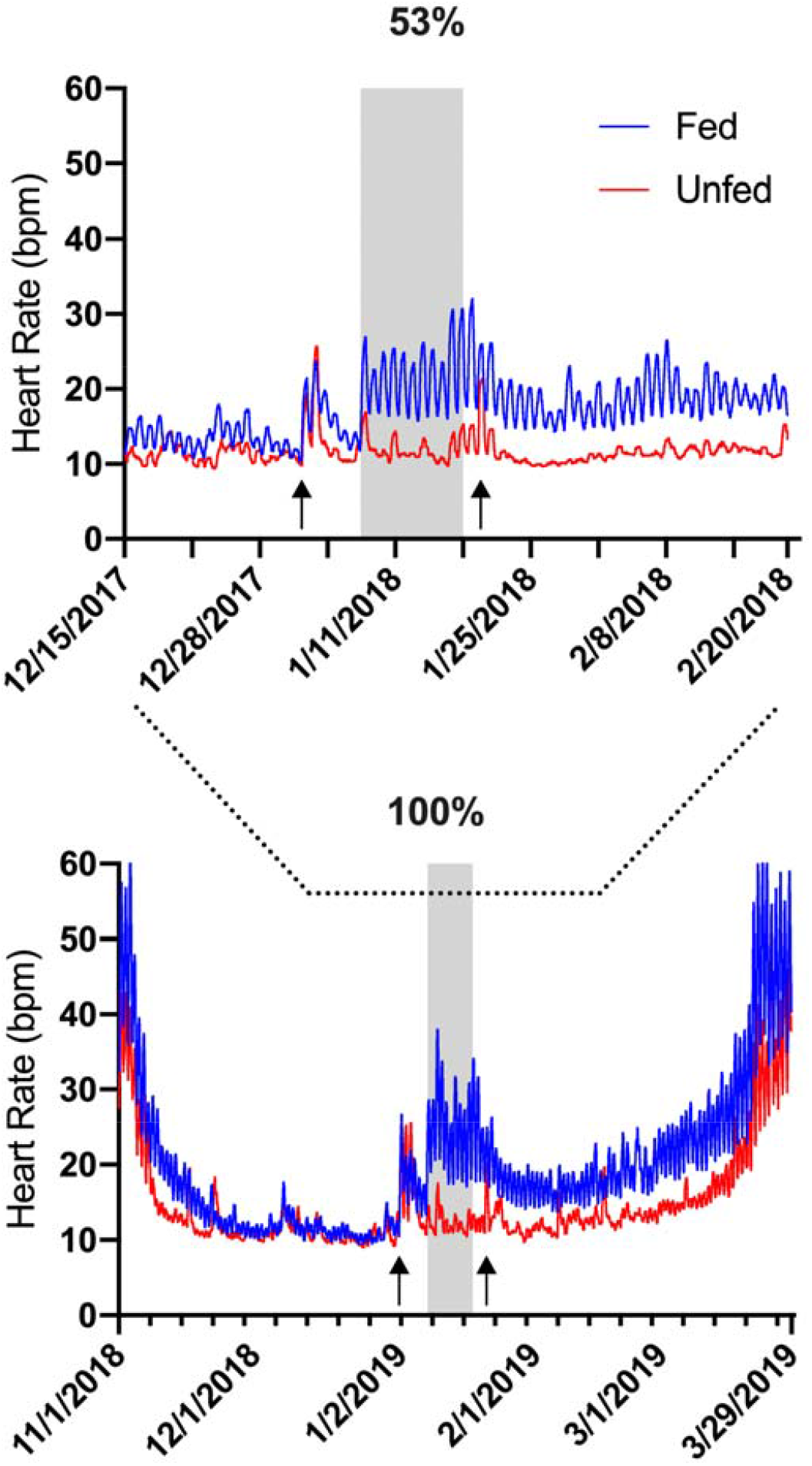
Heart rate data for bears fed two levels of energy replacement (blue, N=4) and a single unfed bear (red). 2 min data are plotted as 200 point moving averages to visualize long-term trends more easily. The same unfed bear (red) is shown in both panels and in two consecutive years. Dashed line between graphs shows the recording period of the first study in relation to the second. Arrows indicate biopsy dates.

Body temperature remained low in all bears and was not significantly affected by feeding (mean±SEM; 53%: Pre-32.91±0.40°C, Post-33.97±0.44°C; 100%: Pre-33.17±0.38°C, Post-34.62±0.96°C; (F(1,12)=4.438; P=0.0569). Body temperature also did not differ between levels of glucose fed (F(1,12)=0.5774, P=0.462).

The strength of the daily heart rate rhythm was low (mean range 14-18%) before glucose feeding but increased significantly in strength to >40% during feeding (Fig. 7); F(2,18)=44.29, P<0.0001. Rhythm strength was then reduced in the 10 days following feeding and was significantly lower in the 100% fed group compared to the 53% group (t(18)=3.090, P=0.0188) (Fig. 7). A significant interaction between experimental phase and feeding level was also observed F(2,18)=6.985, P=0.0057. No effect on rhythm period (peak-to-peak interval) was observed (mean rhythm period = 24.0h).

**FIGURE 7.**
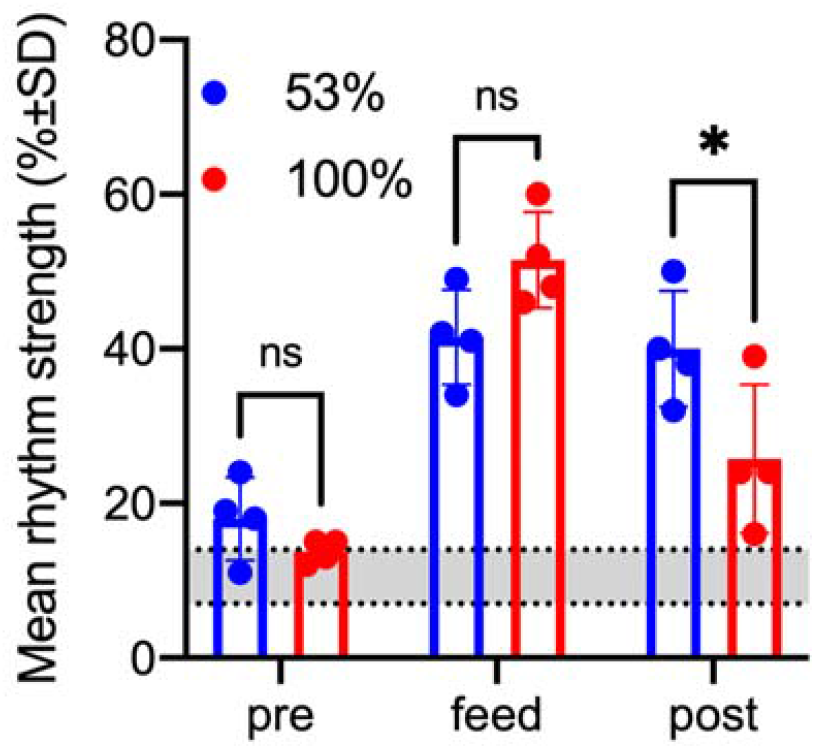
Strength (%) of the daily heart rate rhythm before (pre), during (feed), and after (post) glucose feeding. Gray shading represents the range of heart rate rhythm strength for the unfed bear.

### 5. Cellular energetics

#### a. Mitochondrial respiration

Serum source significantly influenced oxygen consumption under baseline (non-stressed) conditions (F(3,16)=15.70, P<0.0001) (Fig. 8, Fig. S3). Hibernation cells cultured in matching serum (i.e., hibernation, HIBS) exhibited the lowest oxygen consumption rates (0.337 pmol O_2_ min-1 µg protein^-1^) and this rate was 41.8% lower than ACTS (0.478 pmol O_2_ min^-1^ µg^-1^ protein; t(16)=3.152, P=0.0184). Culturing the hibernation cells with serum from fed bears (DEXS, 53%) significantly increased oxygen consumption by 33.6% (0.451 pmol O_2_ min^-1^ µg^-1^ protein; t(16)=2.538, P=0.0434). All hibernation cells cultured in bear serum exhibited lower mitochondrial respiration rates compared to cells cultured with FBS (P≤0.0088). Serum affected oxygen consumption under stressed conditions (F(3,16)=6.503, P=0.0044). Post-hoc analysis revealed that only FBS (P≤0.0166) contributed to the main effect since none of the bear serum treatments differed significantly from one another.

**FIGURE 8.**
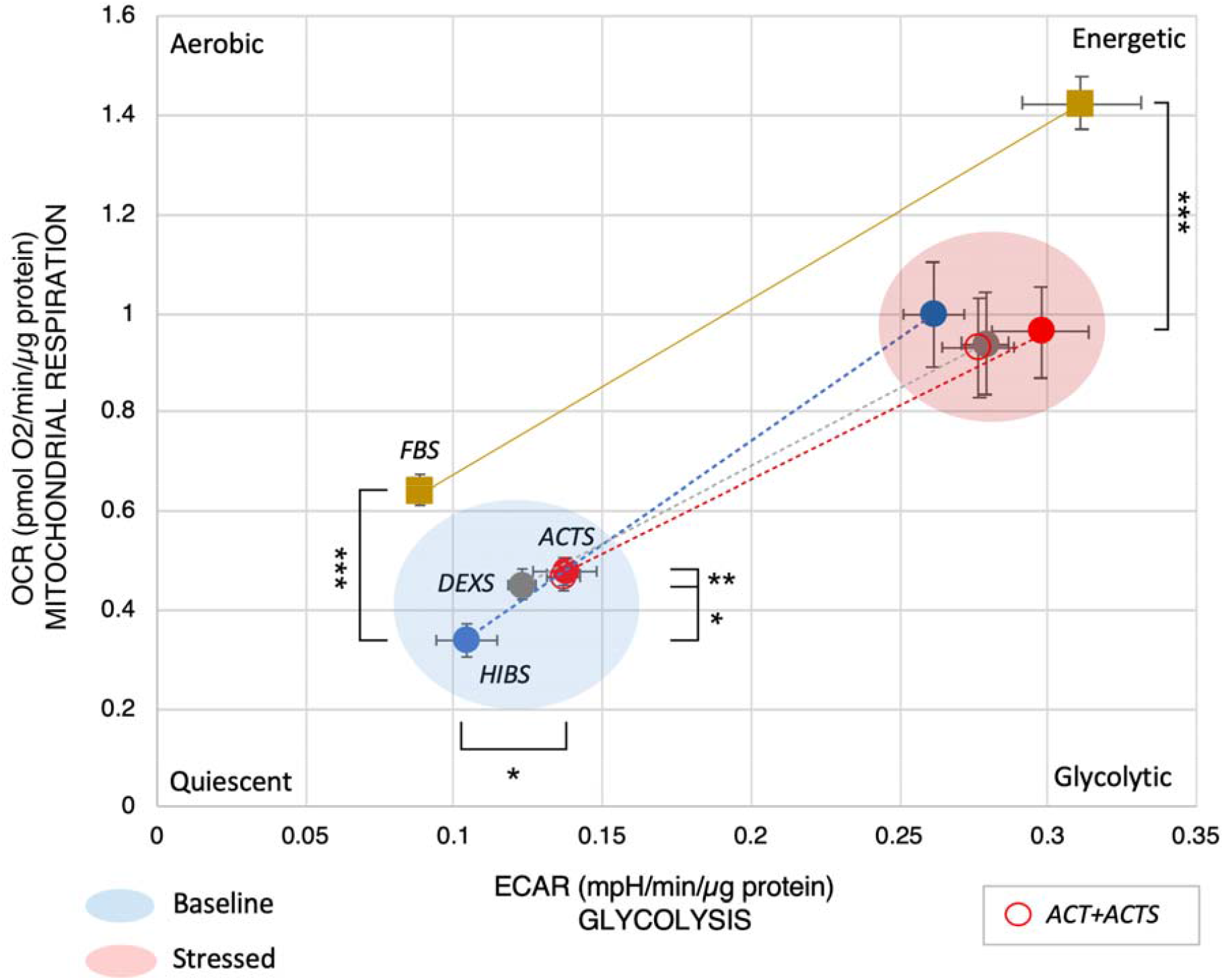
Cell phenotype of bear adipocytes obtained from hibernating bears and cultured in the presence of different serum combinations (10%): HIBS - hibernation serum (prior to feeding); DEXS - serum from fed bears; ACTS - serum from active season bears (Jun-Jul); FBS – fetal bovine serum. Active season cells cultured with active season serum are shown for comparison but were not included in the analysis. * P≤0.05; ** P≤0.01, *** P≤0.001. Results are representative of three separate experiments.

#### b. Glycolytic flux

Hibernation cells cultured with matching (HIBS) serum exhibited the lowest glycolytic flux (Fig. 8, Fig. S3; 0.104 mpH min^-1^ µg^-1^ protein). Under baseline conditions, serum significantly affected medium acidification of hibernation cells (F(3,16)=8.132, P=0.0016). Cells cultured with ACTS exhibited a significantly (33.5%) greater glycolytic flux compared to HIBS (t(16)=3.188, P=0.0283). By contrast, neither FBS nor DEXS caused significant changes in glycolytic flux (i.e. rightward shift) compared to HIBS. No significant differences in stress responses for any cell:serum combination were observed (F(3,16)=2.302, P=0.116).

## DISCUSSION

Hibernation in bears, as in many other species, is a period of energy conservation when food is scarce. The evolution of this process likely involved numerous physiological adaptations for hibernation to be a successful survival strategy (see reviews by (Carey et al., 2003; Geiser, 1998; Geiser, 2013; Melvin and Andrews, 2009). All of the physiological changes that occur during hibernation are eventually reversed once animals exit the den and begin to feed. In an attempt to better understand the processes involved in hibernation, we asked if it was possible to reverse the hibernation state by feeding hibernating bears a single macronutrient, glucose. Three physiologic systems were interrogated in this study: 1) glucose homeostasis, 2) energetics and metabolism and 3) circadian rhythms. The results demonstrate that the systems studied exhibited partial or complete reversal with glucose feeding. This approach could be useful in identifying the critical factors necessary to sustain hibernation in bears and perhaps other species.

We found that blood glucose concentrations at two hours after glucose feeding, irrespective of the amount of glucose fed, returned to levels intermediate to those of hibernation and active seasons suggesting that insulin resistance was partially reversed. This partial reversal is similar to findings in fasted diabetic humans (Cahill et al., 1966). We did not find a significant effect of glucose feeding at the highest level on the insulin:glucose ratio, which has been used as proxy of insulin resistance (Turner et al., 1979) (t(9)=2.003, P=0.0762). This is perhaps not surprising as bears exhibit little evidence of hyperglycemia in hibernation (Rigano et al., 2017; Welinder et al., 2016). A similar lack of significant effect of feeding on insulin:glucose ratios in black bears fed over the winter was observed previously (McCain et al., 2013). Despite the many differences in diet and duration between these two studies, the similar results suggest that factors other than diet are primarily responsible for driving changes in insulin sensitivity. Interestingly, although we found no changes in insulin:glucose ratios in January with feeding, when we performed oGTTs in March (late hibernation) and compared those results with those obtained in June (active season; long-term fed bears) (Fig. S2) we saw large differences in insulin:glucose ratios at baseline (x1000 = March – 0.7, June – 1.2). These results are of similar magnitude and direction to those reported in fasted humans (Cahill et al., 1970). Our results therefore reveal a previously unknown feature of the insulin resistance in hibernating bears, namely, that it progressively increases throughout hibernation. A similar progressive change has been observed in elephant seal pups over several months of fasting (Olmstead et al., 2017).

Elevated circulating FFA have been strongly associated with obesity and insulin resistance in humans (see Review by Boden, 2008)). The reductions in circulating FFA we observed would be consistent with a restoration of insulin sensitivity. However, the suppression was not complete and remained at about 50% greater than active levels. By contrast, the reductions in ketone production were essentially complete and are not unlike those observed in marmots stimulated to feed during hibernation (Tokuyama et al., 1991). These results together highlight the inherent flexibility of metabolic systems in hibernators in response to nutrients. Combined with the observed changes in whole body glucose disposal following single nutrient feeding, this should make the identification of cellular and molecular mediators more straightforward.

Certainly, many effects can be attributed to defects in the insulin signaling pathway (Dresner et al., 1999). Because we previously reported reductions in expression of genes in the insulin signaling pathway occur normally during hibernation (Jansen et al., 2019) we would predict that glucose feeding would reverse those changes. For example, we previously found that expression of the extracellular matrix protein MMP-2, a matrix metalloproteinase was increased in adipose tissue (Jansen et al., 2019). MMP-2 has been linked to elevated FFA and insulin resistance via its ability to cleave the extracellular domain of the insulin receptor (Delano and Schmid-SchoLJNbein, 2008). Additionally, several integrin-related proteins are known to interact with the insulin receptor, such as integrin-linked kinase (ILK) (Williams et al., 2015). We recently found the *ILK* expression was significantly reduced in adipose of hibernating bears (Jansen et al., 2019). Other metabolic pathways involving ketones acting alone or together with fatty acids have also been proposed to confer insulin resistance via disruption of fatty acid oxidation in the mitochondria (Schooneman et al., 2013). Key intermediates in this cascade are the acylcarnitines which are elevated in hibernating bears (Welinder et al., 2016). Acylcarnitines act in the mitochondria via the enzyme carnitine acetyl-CoA transferase (CrAT) (Muoio et al., 2012). *CrAT* gene expression was significantly lower in hibernating bear adipose (Jansen et al., 2019) and in diabetic humans (Muoio et al., 2012). It remains to be determined if changes in the expression of these genes occurs after feeding bears.

Results from our in vitro studies confirmed that adipocytes from fed bears exhibited an enhanced response to insulin. This could not be explained by differences in serum concentrations of glucose or insulin concentrations as these were not different at baseline before or after feeding (Fig. 2). Along the same lines, the failure of hibernating cells to respond to insulin could not be explained by differences in serum insulin concentrations. However, it is possible that longer exposure to glucose could impact the concentrations of insulin and glucose and thereby sensitize the cells to insulin. A more likely explanation is that other serum factors are important for determining insulin sensitivity, metabolism and energetics. In addition to the effects of different sera we also found evidence for cell autonomous effects. For example, glucose uptake was much greater in hibernating cells cultured with active season serum compared to fed cells cultured in fed serum (Fig. 3). This is similar to our previous observations where hibernation cells cultured with active season serum responded more to insulin than active season cells cultured in matching serum (Rigano et al., 2017). It will be important to fully characterize the gene expression changes in cultured adipocytes under similar and contrasting serum conditions to identify the players involved.

It is well established that hibernation is characterized by reductions in acitvity levels and longer torpor bouts in response to the absence of food. Furthermore, the number and duration of torpor bouts can be influenced by diet (Dark, 2005; Frank et al., 2008; Vuarin and Henry, 2014). These effects during hibernation have usually been studied in food-storing hibernators. In several studies higher amounts of polyunsaturated fats (PUFA) in food caches resulted in the shortening of hibernation duration (Munro et al., 2005; Siutz et al., 2017) and supports the hypothesis that increased energy intake shortens hibernation duration. Other studies in bears found no effect of PUFA (Rivet et al., 2017) or were inconclusive in lemurs (Faherty et al., 2017). To our knowledge, a pure carbohydrate has not been adminstererd in hibernation to determine if it can reverse the hibernation state. We found that glucose feeding caused dramatic and prolonged increases in general activity levels, despite being induced by less than two weeks of feeding glucose at a level necessary to offset the lowest predicted cost of hibernation. Along with this we saw a prolonged, roughly 30%, increase in heart rate after feeding. As heart rate is a proxy for metabolic rate, the increase is indicative of increased energy expenditure after feeding.

A controversial aspect of hibernation physiology, namely, the role and importance of circadian rhythms has received relatively little attention in bears (Harlow et al., 2004; Jansen et al., 2016; Körtner and Geiser, 2000; Ruby, 2003; Toien et al., 2015; Ware et al., 2012; Williams et al., 2011). Since bears hibernate at elevated body temperature, questions regarding the integrity and function of circadian rhythms are relevant to our understanding of hibernation. The most striking aspect of the circadian rhythm in hibernating bears is not that it is absent, but that it persists, although at very low amplitude (Jansen et al., 2016). Given the reduction in metabolic rate (up to 75%) during hibernation in bears (Toien et al., 2011; Watts and Cuyler, 1988; Watts and Jonkel, 1988), it is likely that the reduction in circadian amplitude we observed for heart rate prior to feeding is a reflection of decreased energetic demand, nutrient status, or both (Jansen et al., 2016; Ware et al., 2012), although environmental influences cannot be ruled out (Evans et al., 2016). Intriguingly, a role for nutrient status in the operation of the circadian clock has been proposed for numerous species ranging from yeast to mice to maintain the temporal separation of incompatible cellular process (Wang et al., 2015). Thus, the increase in rhythm strength during feeding supports this the hypothesis that circadian clocks are directly responsive to nutrient availability. Our cultured adipocyte model system could lead to new approaches for studying links between circadian rhythms and energetics.

Somewhat surprisingly, the effects of glucose on most parameters were virtually identical regardless of the level of energy replacement. This suggests that there is a ceiling (i.e., 53% of LOMR) beyond which no further increases are possible and that the remainder is due to an active metabolic suppression. However, feeding lower amounts of glucose and/or feeding for longer periods would be needed to confirm this. Alternatively, other dietary factors such as protein or essential fatty acids may be required for full restoration to occur. This seems somewhat unlikely however, as circulating glycerol and ß-hydroxybutyrate concentrations were suppressed to summer active levels with only 53% glucose (Table 1) (Graesli et al., 2015; Rigano et al., 2017). Thus, the most parsimonious explanation for our findings is that fatty acid oxidation was inhibited to a maximum of ∼50% allowing the ingested glucose to become an alternate metabolic fuel, while the remaining ∼50% of fatty acid metabolism was maintained to suppress insulin sensitivity, hence glucose utilization. Altogether, these results reveal a high degree of metabolic flexibility and coordination of physiological processes occurring in hibernating bears.

Metabolic flux analyses revealed a greater than 40% reduction in oxygen consumption and suppression of glycolysis rates in hibernation cells compared to active season cells under season-matching serum conditions. The metabolic suppression occurred at 37°C and thus provides external validation of the proposed independence between temperature and metabolic suppression proposed for bears (Toien et al., 2011). Additionally, the greater distance between basal and stressed levels of mitochondrial respiration of hibernation cells cultured in hibernation serum reveals that hibernation cells have a greater metabolic potential than active season cells or cells from fed bears. This would be predicted if fatty acid oxidation is the primary fuel source as fatty oxidation yields more ATP. Glucose feeding diminished this potential and supports the metabolic switch in fuel use. In summary, it will be possible now to model certain aspects of hibernation “in a dish” for more detailed dissection of the cellular and molecular pathways involved.

## LIMITATIONS AND PROSPECTS

The number of animals used to monitor heart rate was too low to enable statistical analysis. Thus, future studies should include more unfed bears. We also were not able to perform comparisons of glucose uptake and metabolic flux analysis for both levels of glucose feeding. However, given the similarity in the results for all other measures we predict those outcomes would be similar. The potential that the ‘ceiling’ effect proposed was due to factors unrelated to glucose, such as gut distention, seems unlikely for two reasons. First, even unfed bears in our facility drink water (unpublished observations), thus gut distention is a normal part, albeit small, of hibernation in captive bears given ad libitum access to water. Second, we observed increases in metabolic rate in vitro in cells from fed bears. Although we didn’t have both feeding groups to evaluate, this suggest changes are independent of the gut.

We have demonstrated that several features of the physiology of bear hibernation can be reversed with glucose feeding. This was supported by increases in metabolic rate, circadian rhythm strength and the partial restoration of insulin sensitivity. Where applicable, in vitro studies mirrored these findings. Taken together, this ability to study the processes controlling bear hibernation both in vivo and in highly controlled cell cultures provides a new model system to understand hibernation.

## Acknowledgements

We are grateful to Tim Laske at Medtronic (Minneapolis, MN) for the gift of Reveal LINQ monitors used in the current study, to the many volunteers working at the WSU Bear Center and to Jessie McCleary, Nina Woodford and Gaylynn Clyde of the WSU Office of the Campus Veterinarian.

## Supplementary Material

Supplementary figures are included

## SUPPLEMENTAL FIGURES

**FIGURE S1.**
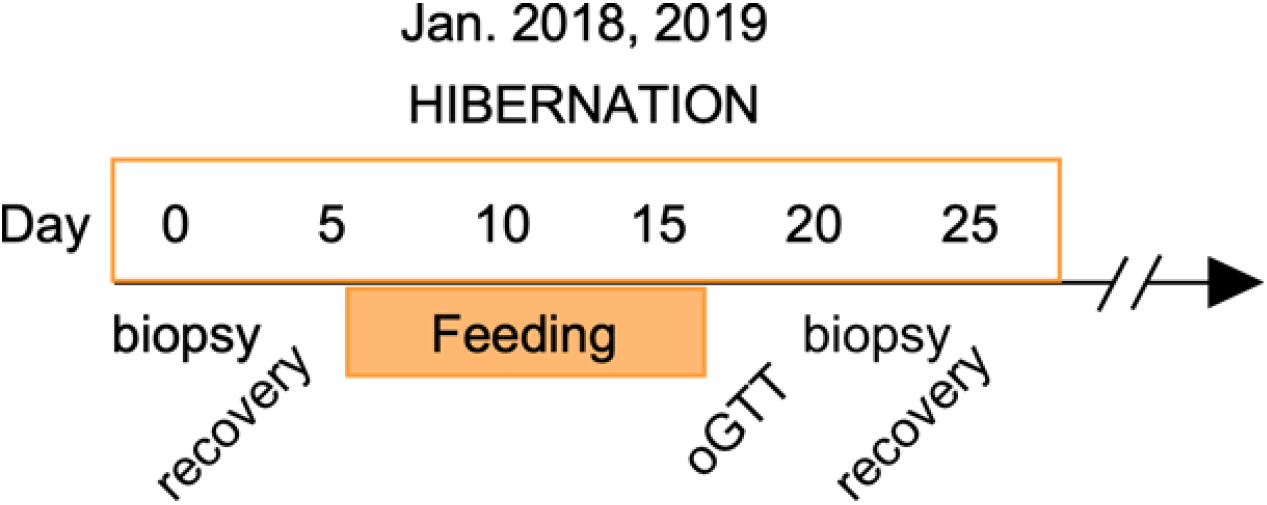
Timeline of procedures for feeding study.

**FIGURE S2.**
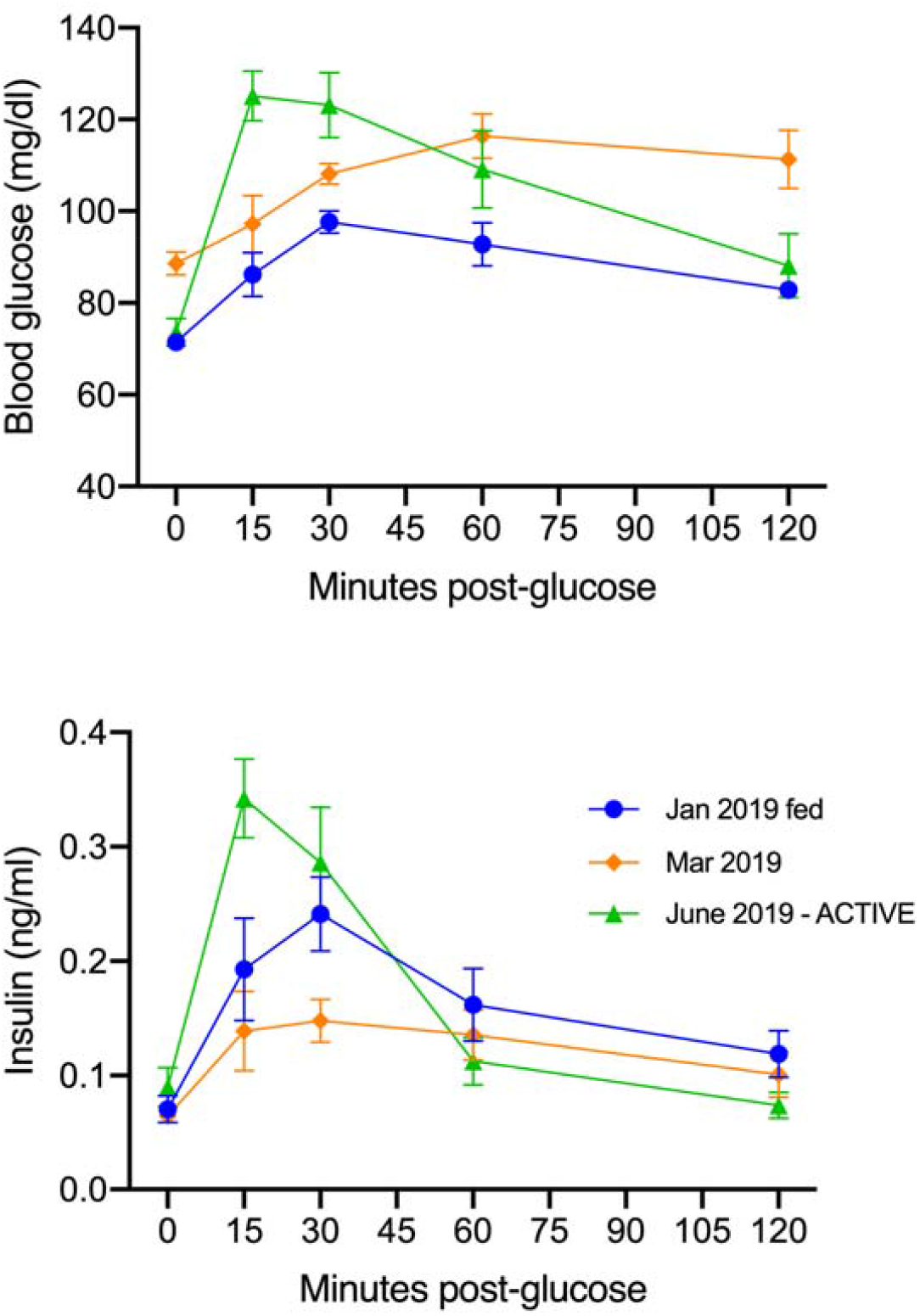
Oral glucose tolerance test results for bears sampled in late hibernation (March 2019) compared to January (2019 post-feeding) and June 2019 (active season). January and June data are the same shown in Fig. 2 of the main text. Values are means ± SEM.

**FIGURE S3.**
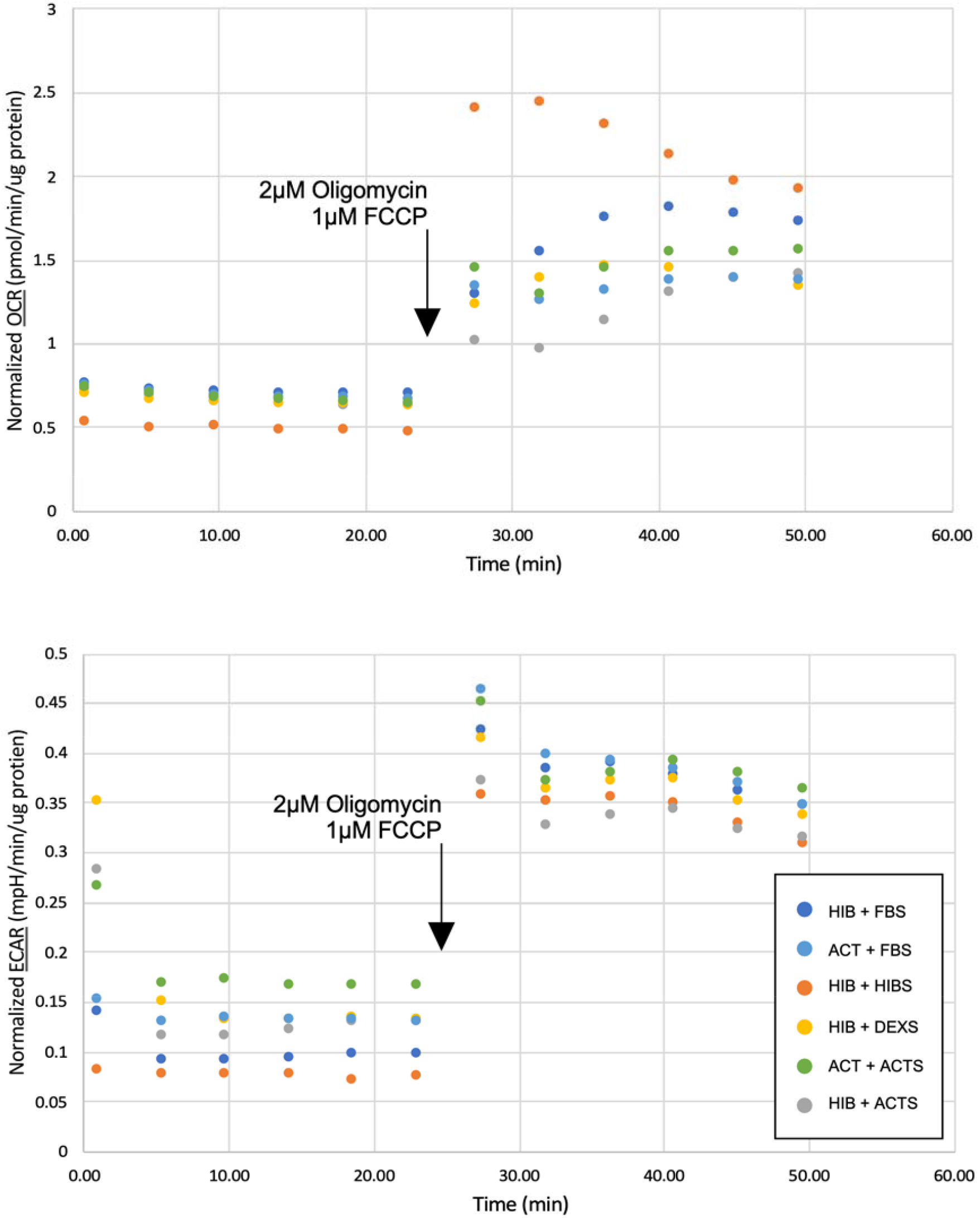
Representative traces of OCR and ECAR measurements in the Seahorse XFp flux analyzer (Agilent, San Diego, CA) from an adipocyte culture under different serum conditions (see Methods for details). All results are from the same bear.

## Notes

### Competing Interest Statement

The authors have declared no competing interest.

